# Is what you see what you get? The relationship between field observed and actual aphid parasitism rates in canola crops

**DOI:** 10.1101/2021.04.12.439466

**Authors:** Samantha Ward, Paul A. Umina, Hazel Parry, Amber Balfour-Cunningham, Xuan Cheng, Thomas Heddle, Joanne C. Holloway, Caitlin Langley, Dustin Severtson, Maarten Van Helden, Ary A. Hoffmann

## Abstract

**BACKGROUND:** Estimating parasitoid abundance in the field can be difficult, even more so when attempting to quantify parasitism rates and the ecosystem service of biological control that parasitoids can provide. To understand how ‘observed’ parasitism rates (in-field mummy counts) of the green peach aphid, *Myzus persicae* (Sulzer) (Homoptera: Aphididae) translate to ‘actual’ parasitism rates (laboratory-reared parasitoid counts), field work was undertaken in Australian canola fields over a growing season. Parasitoids were reared within a controlled laboratory setting.

**RESULTS:** Total observed and actual parasitism rates of *M. persicae* varied considerably across regions, but less so on a field level. Overall, actual parasitism was on average 2.4 times higher than that observed in the field, with rates an average of 4-fold higher in South Australia. As crop growth stage progressed, the percentage of mummies observed increased. Percentage of parasitoids reared also increased with crop growth stage, averaging 3.4% during flowering and reaching 14.4% during podding/senescing. Although there was a greater diversity of reared parasitoid species at later crop growth stages, actual parasitism rate was unaffected by parasitoid species. *Diaeretiella rapae* was the most commonly reared parasitoid, increasing in abundance with crop growth stage.

**CONCLUSION:** These findings indicate that mummy counts alone do not provide a clear representation of parasitism within fields.

## 1. Introduction

The green peach aphid, *Myzus persicae* (Sulzer) (Homoptera: Aphididae), is an important agricultural pest worldwide and was first recorded in Australia in New South Wales (NSW), in 1910^1^. Globally, at least 50 parasitoid species have been reported to attack *M. persicae*^2^, as listed by van Emden et al. (1969)^3^. In Australia, although many of these natural enemies are present, little research has been undertaken to investigate parasitoid species composition, the use of parasitoids as biological control agents of *M. persicae*, the thresholds required to suppress *M. persicae* populations, and the effect of seasonal changes on naturally occurring populations and parasitism rates.

Estimating parasitoid abundance in the field can be difficult, even more so when attempting to understand in-field parasitism rates. Often a parasitized aphid can be identified visually; parasitoids protect themselves by pupating within the eaten-out husk of their aphid host after cementing it to the substrate. This husk is usually referred to as a ‘mummy’^4^. Within Australia, growers and agronomists may use in-field mummy counts as an indicator of aphid parasitoid activity. Based on their findings, usually as part of an Integrated Pest Management (IPM) program, growers judge whether a field should be chemically treated to reduce pest aphid abundance, or whether parasitoid numbers are sufficient to control the aphid populations without the need for spraying. Furthermore, growers can opt to invest in (often more expensive) selective insecticides, as opposed to (often cheaper) broad-spectrum insecticides, to protect the aphid parasitoids if they are thought to be providing control of aphids longer term. Unfortunately, however, these crude counts using mummies can prove inaccurate.

Powell et al. (1996)^5^ suggested that aphid mummy counts should be combined with other methods to estimate parasitoid abundance, because the sampling method alone could lead to misleading results. Mummified aphids may not be distinguishable from unparasitized individuals for several days, resulting in an underestimate of total parasitism. Mummified aphids change into a golden yellow colour, the recognisable trait of mummification, approximately 72 hours after oviposition at 23°C^6^. Based on visual monitoring alone, any *M. persicae* mummified at this temperature in the field <72 hours prior could be incorrectly presumed to be unparasitized. Uncertainty is likely to be higher as temperature drops and/or becomes more variable. Another reason is that strong wind or heavy rain can dislodge mummies from plant leaves^5^.

In the field, regardless of host species, many parasitoids are likely to die during development, with Nielsen and Hajek (2005)^7^ noting an overall rate of 56% non-emergence across hosts. Dissection of mummies that failed to rear parasitoids revealed the parasitoid to be either dead or diapausing^8^. Walton et al. (1990)^8^ found that mummy counts generally underestimated parasitism levels, when compared to live rearing or electrophoresis. Giles et al. (2003)^9^ also noted a weak relationship between the proportions of tillers with mummies against proportions of parasitised aphids in wheat and these authors suggested that simultaneous aphid and parasitoid sampling is required to determine whether chemical applications are required^9^. These findings reinforce the notion that mummy counts alone are unlikely to provide sufficient information to accurately estimate parasitoid density.

The aims of this paper were to understand how observed parasitism rates of *M. persicae* relate to the actual parasitism rates in canola fields (*Brassica napus* L.) throughout Australia, and whether these rates varied through the growing season as well as across regions. We also investigated whether these rates varied spatially across a field, in relation to the distance from the field edge, and whether the composition of aphid parasitoids affected estimates of observed and actual parasitism rates.

## 2. Materials and Methods

### 2.1. Site selection

In 2019, 10-13 seed-treated canola fields, at least 1 km from one another, were surveyed for *M. persicae* in each of the following states: New South Wales (NSW), South Australia (SA), Victoria (VIC), and Western Australia (WA). Site visits were conducted from June/July 2019 every four weeks. Once aphids were detected, only five fields with *M. persicae* in SA and WA, six in NSW, and seven in VIC, continued to be sampled until the end of the season (as defined by the time of windrowing of the crop) (Fig. 1). Selection of these fields was based on insecticide usage, with preference given to those with no sprays.

**Figure 1:**
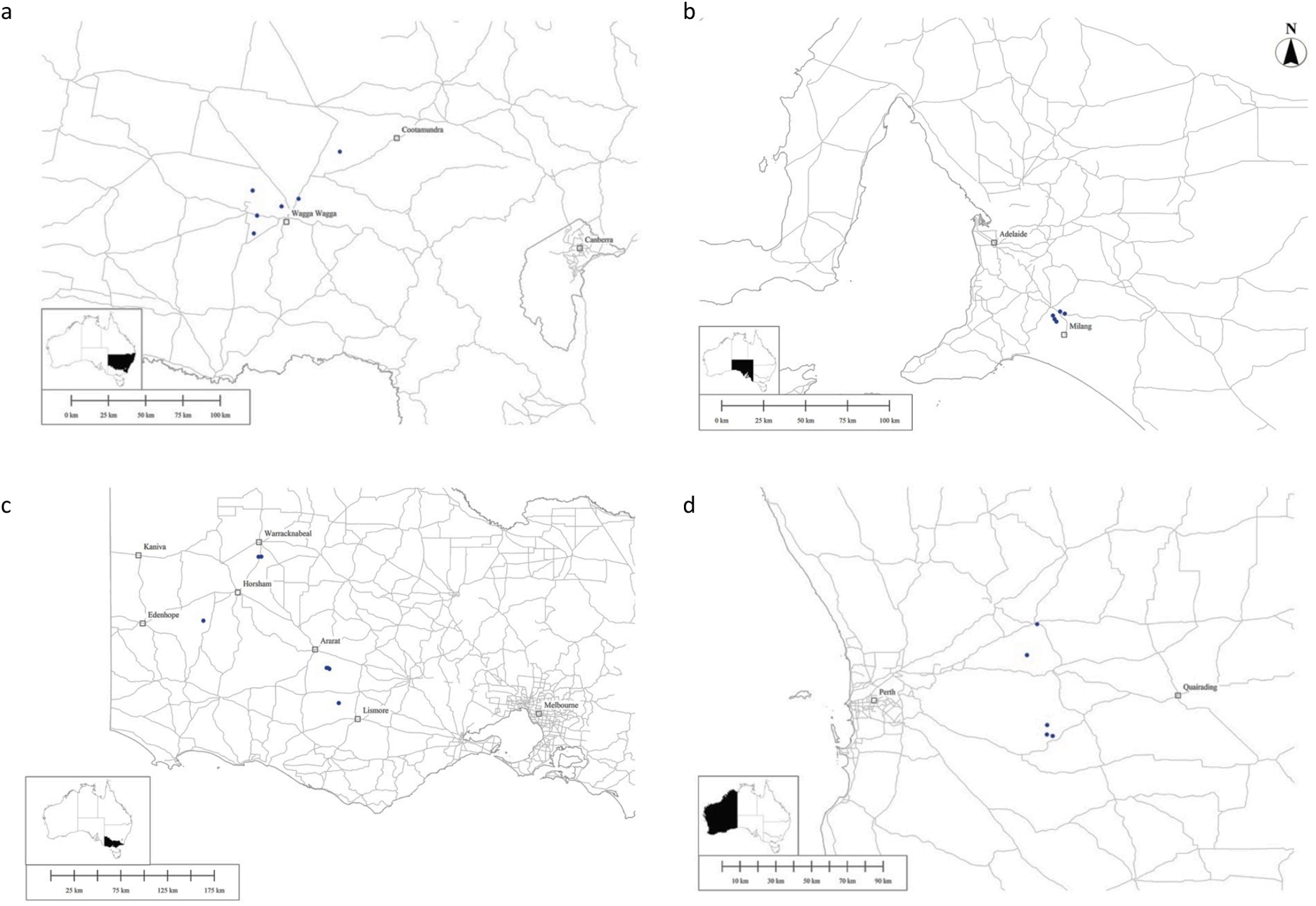
Map of field sites in each state: a) NSW, b) SA, c) VIC, and d) WA (inset maps depict states on Australian map)^10,11^.

### 2.2. Aphid collections

Plants within fields were sampled directly by hand for the presence of aphids and mummies, at sampling points at least 30 m from one another. Plant condition was categorised as ‘unstressed’, where plants looked healthy and turgid, or ‘stressed’ where plants were wilting, had a yellowing of or dull colour to leaves, were stunted, patchy, or had white feeding marks on the leaves, or a combination of the above. Stressed plants were targeted, and searches began at the edge of the crop, moving into the field, in a zigzag formation, reaching a distance of >100 m from the field edge. Plants were searched with a focus on the underside of the lower leaves of canola plants due to this being the most common location for *M. persicae*^12^. *Myzus persicae* was targeted and prioritised over other aphid species when present, however the presence of cabbage aphid (*Brevicoryne brassicae* L.) and turnip aphid (*Lipaphis erysimi* (Kaltenbach)), was noted. Each sampling point was searched at random for *M. persicae* aphids and mummies for one minute, or until the combined count of these reached 50 individuals, whichever occurred first. Sampling points were inspected until eight sites positive for *M. persicae* were logged. If *M. persicae* was absent, a sampling point was recorded, before another point was selected for inspection, until a maximum of 24 sampling points had been inspected at each field.

Direct sampling of aphids was undertaken with the help of a mobile software application developed for this project by Andy Hulthen (CSIRO) for electronic data collection using the Open Data Kit described in Hartung et al. (2010)^13^. The application accessed the in-built GPS and location-based services of the mobile phone as well as a barcode-reading capability for recording individual samples labelled with barcodes. The application was used to electronically record field-derived (and subsequently lab-derived) data. The data were uploaded directly to a database in the Cloud (when in mobile internet range). Collectors followed prompts on the field mobile application, noting several variables, such as crop growth stage, plant condition, alate/nymph presence, and natural enemy presence (see Tables S1, S2).

Aphids and mummies were kept on their respective leaves and stored, along with paper towelling and leaves lacking aphids, in a sealed container. Each sampling container was labelled with a barcode that corresponded with the phone application record. Samples were kept refrigerated or on ice for transport to the laboratory.

### 2.3. Laboratory work

Parasitoids were reared from both field-collected aphids and mummies as described below and then identified to species. For material collected from NSW fields, data was not recorded on whether a parasitoid emerged from the mummies or the aphids collected from the field. For the other states, these two groups were recorded separately. Consequently, NSW data was omitted from some analyses.

#### 2.3.1. Rearing from field collected mummies

All mummies collected from a sampling point were placed into a petri dish lined with moist filter paper. The underside of each petri dish was labelled with the corresponding barcode and placed within a controlled temperature (CT) cabinet maintained at 20°C and a 16L:8D photoperiod. For two weeks, petri dishes were checked every second day for parasitoid emergence. Once a parasitoid emerged, it was stored in 80 % ethanol, and labelled with a corresponding barcode. Parasitoids reared from the same sampling point were stored together.

#### 2.3.2. Rearing from field collected (non-mummified) aphids

Petri dishes were made up with a 1 % agar solution, within which 2-3 cotyledons of canola or sprouting radish (*Raphanus raphanistrum* subsp. *sativus* L.) were inserted. The lid was lined with filter paper. Aphids from the same sampling point were placed onto the cotyledons, unless numbers were very high (∼50 or above), in which case multiple petri dishes were used. Each dish was labelled on the underside with the corresponding barcode and placed within the CT cabinet maintained at 20°C and a 16L:8D photoperiod. Leaves were changed weekly, or if limp or discoloured, or if fungus began to appear. Any mummies that developed were removed and placed in a separate petri dish. These mummies were checked every second day for parasitoid emergence for two weeks from collection. Once the parasitoid emerged it was stored in 80 % ethanol separate to the mummy case. At the end of the two weeks, all aphids that remained unparasitized were stored in 80 % ethanol. For NSW, parasitoids produced from mummies and non-mummified aphids were combined within the same tubes with 80 % ethanol. All parasitoids reared from *M. persicae* were stored at 4°C and identified morphologically to species level, using keys by Rakhshani et al. (2012,2015)^14,15^.

#### 2.3.3. DNA extraction methods & PCR amplification

Parasitoid DNA was extracted non-destructively using a modified Chelex® extraction method, adapted from Walsh et al. (1991)^16^, as detailed in Carew et al. (2003)^17^. An individual parasitoid was placed within a micro-centrifuge tube, along with 3 µl of Proteinase K (20mg/ml) and 70 µl of 5 % Chelex® solution, before being incubated in a water bath. PCR was undertaken using a 10 % dilution of the DNA extractions, amplifying the samples with the “universal” arthropod primer pair LCO1490/HCO2198^18^. Reactions contained a final concentration of 1x Standard Taq Reaction Buffer (New England Biolabs, Massachusetts, USA), 2.5 mM MgCl_2_, 0.5 µM each primer, 0.2 mM dNTPs, 2.4 U IMMOLASE DNA Polymerase (Bioline, London, UK) and 3 µL diluted DNA, in a reaction volume of 30 µL. Amplicons were sent to the Australian Genome Research Facility (AGRF) for sequencing, before forward and reverse sequences were assembled and trimmed using Geneious version 9.1.8 (https://www.geneious.com). Sequences were identified using the Genbank database (http://www.ncbi.nlm.nih.gov) and cross-referenced with the Barcode of Life Data System database (BOLD; http://www.barcodinglife.org^19^.

### 2.4. Data analyses

Parasitism rates were calculated in two ways. The first was the ‘observed parasitism’ rate computed as the number of mummies observed in the field compared to the total number of aphids and mummies collected. The second was the ‘actual parasitism’ rate, defined as the number of parasitoids that were reared divided by the number of aphids and mummies that were sourced. The ratio of parasitoids reared from mummies collected from the field (‘field mummies’) over the number reared from aphids collected from the field that subsequently became mummified (‘field aphids’) was also investigated in order to assess the extent to which field mummy counts might underestimate parasitoid impact.

General Linear Models (GLMs), with fields as a factor nested within states, were undertaken to analyse the effects of state, field, crop growth stage, and crop stress on the variables. Count data were log transformed (ln(X+1)) prior to analysis as they were not normally distributed, while proportion data were logit transformed as recommended by Warton et al. (2011)^20^. We used this approach rather than treating the presence of alates, mummies and parasitoids as binomial variables because of the uneven sampling and patchy distribution of aphids, and potentially parasitism, across fields. Note that proportions were based on the presence of at least 18 aphids/alates/mummies. Proportions of parasitoids reared were based on the presence of at least one aphid. Crop growth stage was considered a factor due to the categorical nature of the variable, and stages prior to flowering were excluded from analysis due to the rarity of aphids. The difference between the number of parasitoids reared from field mummies over the number reared from field aphids was also investigated in order to assess the extent to which field mummy counts might underestimate parasitoid impact. Post hoc Tukey’s multiple comparison tests were undertaken to determine which means differed.

To explore the relationships between abundance of *M. persicae*, proportion of mummies from total aphids sampled, and proportion of parasitoids reared against distance from field edge, GLMs were performed on data with distance from edge as a covariate, after collating data across all growth stages. For these analyses, any fields with fewer than 10 aphids or parasitoids reared were removed.

To assess associations between the measures of parasitism as well as aphid counts, Spearman rank correlations were computed at the field level, per crop growth stage. When comparing *M. persicae* abundance with mummy counts/proportions, data were excluded when parasitism was absent. We also tested whether observed and actual parasitism differed across all spatial data points where parasitism was recorded (pooled across crop growth stages) by using a Sign test.

For the analysis of parasitoid species composition, parasitoid counts were summed across sampling points within a field before multiple response permutation procedures (MRPPs) were used to investigate the effects of state, crop stress, crop growth stage, and field on parasitoid species composition, with Euclidean distance as a similarity measure. Paired t-tests were undertaken to compare the proportion of *D. rapae* to the other parasitoids when reared from field aphids versus field mummies.

All analyses were conducted using Minitab version 19.1.0.0^21^, with the exception of the MRPPs, which were performed in R version 4.0.1^22^, using RStudio version 1.3.959^23^.

## 3. Results

### 3.1. Myzus persicae *abundance at a field level and the proportion of alates, mummies and reared parasitoids*

During 2019, 11246 non-mummified *M. persicae* were collected, with 3578 from sampling points in NSW, 4218 in SA, 585 in VIC, and 2865 in WA (Fig. S1a). In all states, *Myzus persicae* presence varied over time, with no aphids found during the seedling crop growth stage, and very few during the vegetative stage. Aphids were first recorded in low numbers in VIC in early August and in the other states in mid-July. The non-mummified *M. persicae* counts were significantly different between states, but not between fields (Table 1), with the fewest mean aphids per field found in VIC (Figs. 2a, S1a). There was no difference in aphid numbers collected per field at the three later crop growth stages (flowering, flowering/podding, podding/senescence) and no impact of crop stress on numbers (Table 1). In NSW, most sampling was undertaken during the flowering crop growth stage (43 % of visits), in VIC during the flowering/podding stage (44 %), in WA during the flowering/podding and podding/senescing stages (36 %), and in SA during the podding/senescing crop growth stage (41 %).

**Table 1:** Results of GLMs testing the effects of state, field, crop growth stage and crop stress on *M. persicae* numbers, and proportions of alates, mummies, and reared parasitoids considered at the field level.

| Organism & measurement | Factor | MS | F<sub>(df1, df2)</sub> | P |
| --- | --- | --- | --- | --- |
| **Mean *Myzus persicae* abundance** | State | 5.811 | 11.64<sub>(3, 19)</sub> | **<0.001** |
|  | Field (nested within state) | 0.563 | 1.13<sub>(19, 35)</sub> | 0.369 |
|  | Crop growth stage | 1.180 | 2.36<sub>(2, 35)</sub> | 0.109 |
|  | Crop stress | 0.864 | 1.73<sub>(1, 35)</sub> | 0.197 |
|  | Error | 0.499 |  |  |
| **Proportion of alates** | State | 5.158 | 4.84<sub>(3, 17)</sub> | **0.009** |
|  | Field (nested within state) | 0.583 | 0.55<sub>(17, 25)</sub> | 0.899 |
|  | Crop growth stage | 1.545 | 1.45<sub>(2, 25)</sub> | 0.253 |
|  | Crop stress | 1.177 | 1.11<sub>(1, 25)</sub> | 0.303 |
|  | Error | 1.065 |  |  |
| **Proportion of mummies** | State | 5.599 | 3.85<sub>(3, 19)</sub> | **0.019** |
|  | Field (nested within state) | 2.311 | 1.59<sub>(19, 30)</sub> | 0.125 |
|  | Crop growth stage | 41.643 | 28.61<sub>(2, 30)</sub> | **<0.001** |
|  | Crop stress | 0.003 | 0.00<sub>(1, 30)</sub> | 0.962 |
|  | Error | 1.456 |  |  |
| **Proportion of reared parasitoids (exc. NSW)** | State | 2.088 | 1.78<sub>(2, 14)</sub> | 0.192 |
|  | Field (nested within state) | 1.212 | 1.03<sub>(14, 23)</sub> | 0.459 |
|  | Crop growth stage | 6.461 | 5.50<sub>(2, 23)</sub> | **0.011** |
|  | Crop stress | 4.903 | 4.17<sub>(1, 23)</sub> | 0.053 |
|  | Error | 1.175 |  |  |

| Organism & measurement | Factor | MS | F(df1, df2) | P |
| --- | --- | --- | --- | --- |
| **Mean *Myzus persicae* abundance** | State | 5.811 | 11.64<sub>(3, 19)</sub> | **<0.001** |
|  | Field (nested within state) | 0.563 | 1.13<sub>(19, 35)</sub> | 0.369 |
|  | Crop growth stage | 1.180 | 2.36<sub>(2, 35)</sub> | 0.109 |
|  | Crop stress | 0.864 | 1.73<sub>(1, 35)</sub> | 0.197 |
|  | Error | 0.499 |  |  |
| **Proportion of alates** | State | 5.158 | 4.84<sub>(3, 17)</sub> | **0.009** |
|  | Field (nested within state) | 0.583 | 0.55<sub>(17, 25)</sub> | 0.899 |
|  | Crop growth stage | 1.545 | 1.45<sub>(2, 25)</sub> | 0.253 |
|  | Crop stress | 1.177 | 1.11<sub>(1, 25)</sub> | 0.303 |
|  | Error | 1.065 |  |  |
| **Proportion of mummies** | State | 5.599 | 3.85<sub>(3, 19)</sub> | **0.019** |
|  | Field (nested within state) | 2.311 | 1.59<sub>(19, 30)</sub> | 0.125 |
|  | Crop growth stage | 41.643 | 28.61<sub>(2, 30)</sub> | **<0.001** |
|  | Crop stress | 0.003 | 0.00<sub>(1, 30)</sub> | 0.962 |
|  | Error | 1.456 |  |  |
| **Proportion of reared parasitoids (exc. NSW)** | State | 2.088 | 1.78<sub>(2, 14)</sub> | 0.192 |
|  | Field (nested within state) | 1.212 | 1.03<sub>(14, 23)</sub> | 0.459 |
|  | Crop growth stage | 6.461 | 5.50<sub>(2, 23)</sub> | **0.011** |
|  | Crop stress | 4.903 | 4.17<sub>(1, 23)</sub> | 0.053 |
|  | Error | 1.175 |  |  |

**Figure 2:**
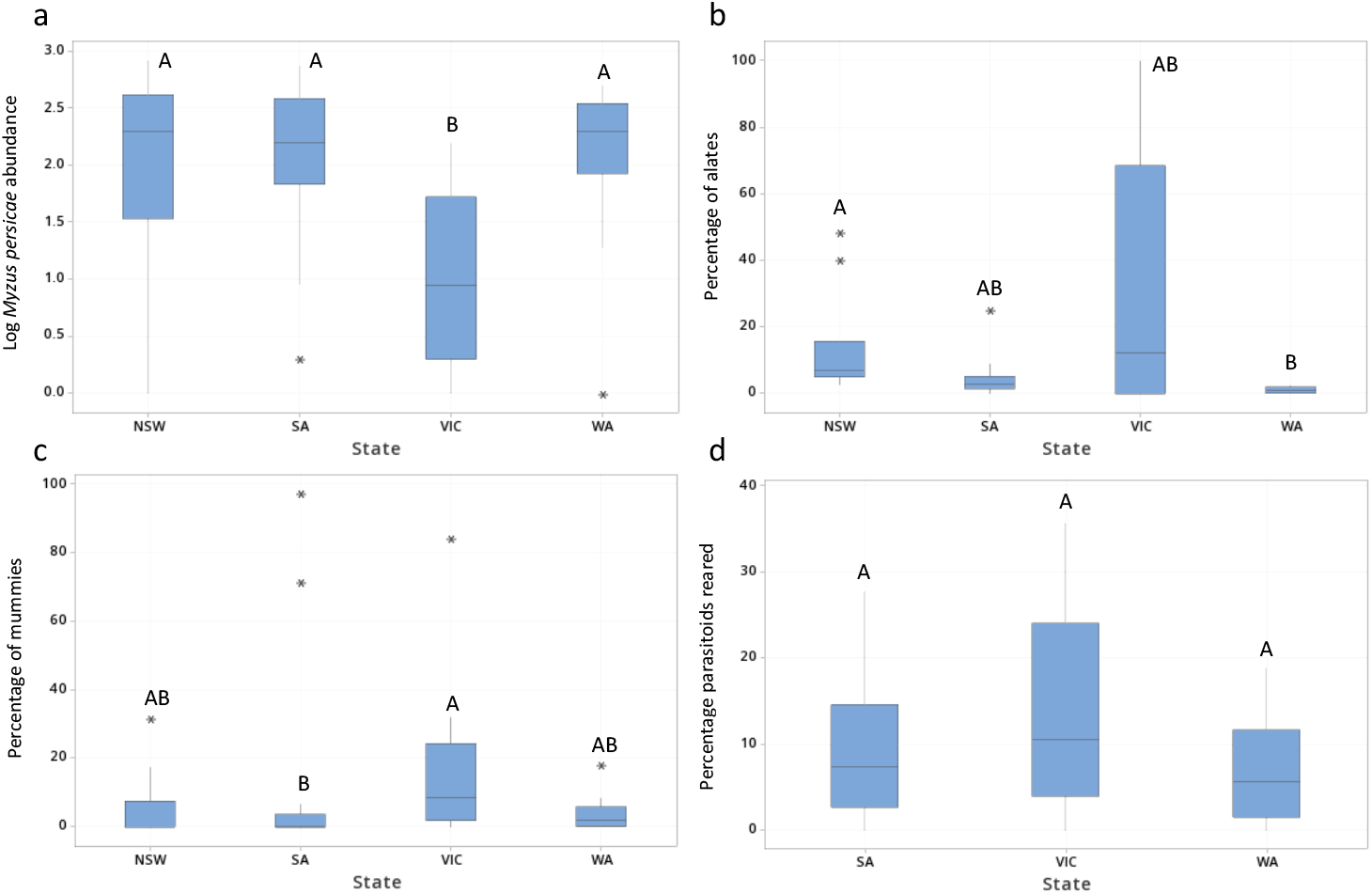
Box plots showing by state, a) *M. persicae* abundance, and the percentage of b) alates, c) mummies and d) reared parasitoids on fields excluding those in NSW. [Outliers are indicated by single asterisks. Different letters indicate significant differences between states in post hoc tests].

The proportion of *M. persicae* alates from total aphids sampled also differed significantly by state (Table 1), with the proportion of alates significantly higher in NSW than in WA (Fig. 2b). The proportion of alates was not affected by field, crop growth stage or crop stress (Table 1).

During 2019, 515 mummies were collected, with 145 in NSW (4 % of total *M. persicae* collected in this state), 148 in SA (4 %), 52 in VIC (9 %) and 170 in WA (6 %) (Fig. S1b). No mummies were found in NSW, SA, and WA until August, and September in VIC. For the proportion of aphids that were collected as mummies, there was a significant effect of state (Table 1), with significantly higher proportions found in VIC than in SA (Fig. 2c). Crop growth stage also significantly affected the proportion of mummies collected (Fig. 3a; Table 1), with a significantly higher proportion of mummies collected during the podding/senescing stage, than during the flowering and flowering/podding stages (Fig 3a). No mummies were found during the seedling stage and only one during the vegetative stage (from NSW). The increase in the proportion of mummies at later crop growth stages reflects an increase in field parasitism estimates consistent across all the states except in NSW (Fig. 3a). The proportion of mummies from total aphids collected was not affected by field or by crop stress (Table 1).

**Figure 3:**
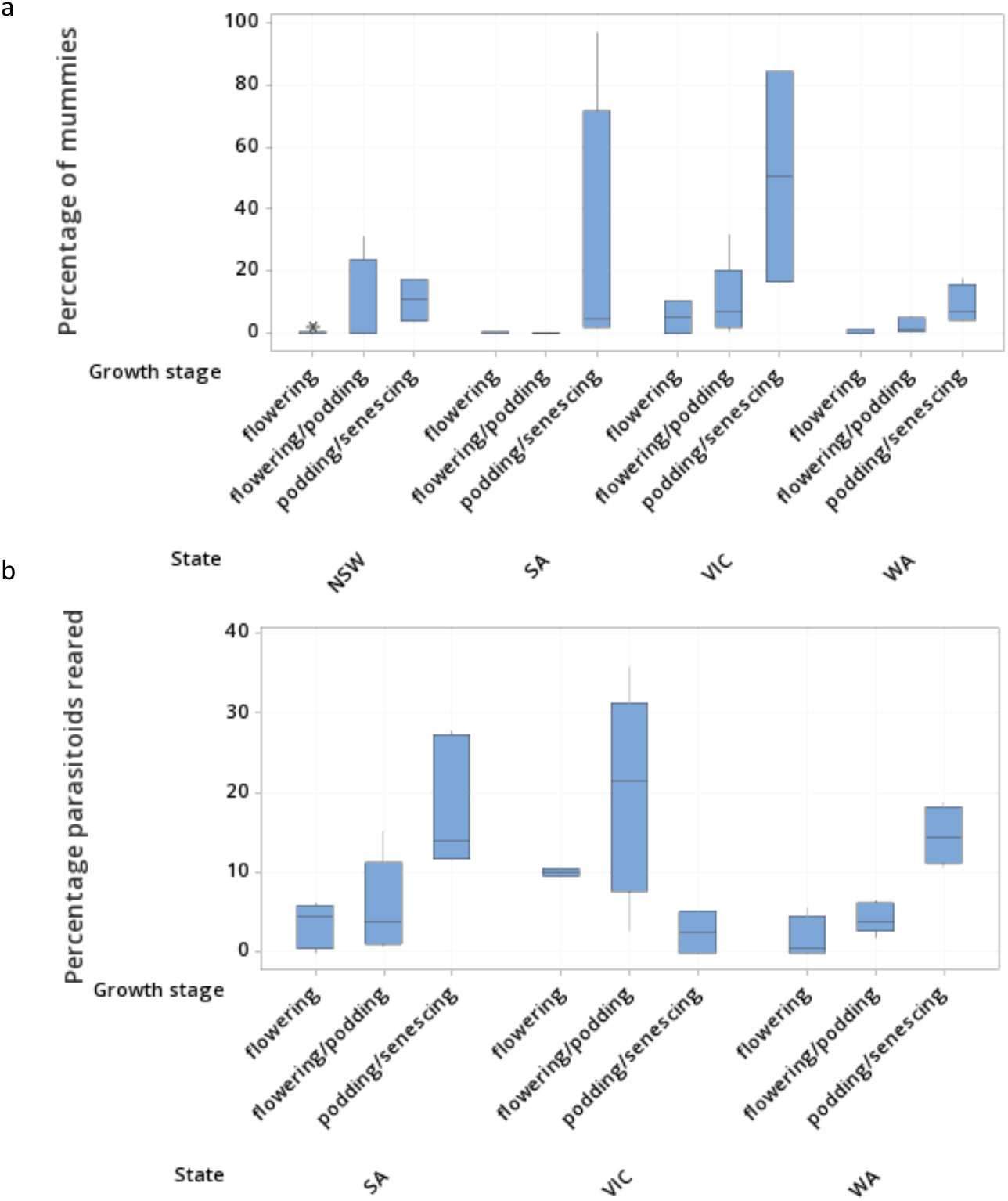
Box plots displaying effects of crop growth stage on a) percentage of samples collected as mummies per state and b) the percentage of samples collected which reared parasitoids per state. [Outliers are indicated by single asterisks. A post hoc test found the percentage of mummies from total aphids collected was significantly higher during the podding/senescing crop growth stage than during flowering or flowering/podding]. A post hoc test found the percentage of mummies from total aphids collected was significantly higher during the podding/senescing crop growth stage than during flowering].

Parasitism as measured by reared parasitoids showed significant effects of crop growth stage, but no effect of state, field, or crop stress (Table 1). As was the case for mummification rates, the proportion of reared parasitoids from total aphids and mummies collected increased with crop growth stage in both SA and WA, although this was not the case in VIC (Fig. 3b).

### 3.2. Comparison of observed and actual rates of parasitism

Across all fields, 515 mummies were sampled during this study, while 1221 parasitoids were successfully reared in the laboratory, suggesting that the total actual parasitism rate was overall 237 % higher than that estimated in the field based on mummy counts (observed parasitism rate). At the state level, 145 mummies were collected, and 233 parasitoids were reared in NSW, presenting an actual parasitism rate that was 161 % of the observed parasitism rate; similarly, 148 mummies were collected, and 606 parasitoids reared in SA (409%), 52 mummies were collected, and 92 parasitoids were reared in VIC (177 %), and 170 mummies were collected, and 290 parasitoids were reared in WA (171 %).

Comparing observed and actual parasitism rates at the field level, of the 61 fields sampled (separated by time points), 27 had higher actual parasitism rates than observed parasitism rates (44 %), 15 had the same parasitism rates (25 %), and 19 had less actual parasitism than observed (31 %; due to failed rearings). We also looked at observed and actual parasitism rates at the level of individual sampling points in a field. When combined over time, of the 88 sampling points that were repeatedly sampled, where parasitism was observed as either field mummy counts or resulted in reared parasitoids in SA, VIC and WA, 83 % (73) had higher actual parasitism than observed parasitism, 10 % (9) had the same parasitism rates, and 7 % (6) had less actual parasitism than observed. These comparisons support the notion that actual parasitism was higher than observed parasitism, particularly at the finer scale of sampling point (Sign test, p**<0.001**). On a field level, when parasitism occurred, the difference in observed and actual parasitism rate (number of parasitoids reared subtracted by mummy count) did not vary across state, field, crop growth stages or crop stress (GLM: state MS=0.364, F_2,12_ =0.97; p=0.397; field MS=0.050, F_12,18_ = 0.13; p=1.000; crop growth stage MS=0.234, F_2,18_=1.18, p=0.546; crop stress MS=0.060, F_1,18_=0.16, p=0.695; error MS=0.374) (Fig. 4).

**Figure 4:**
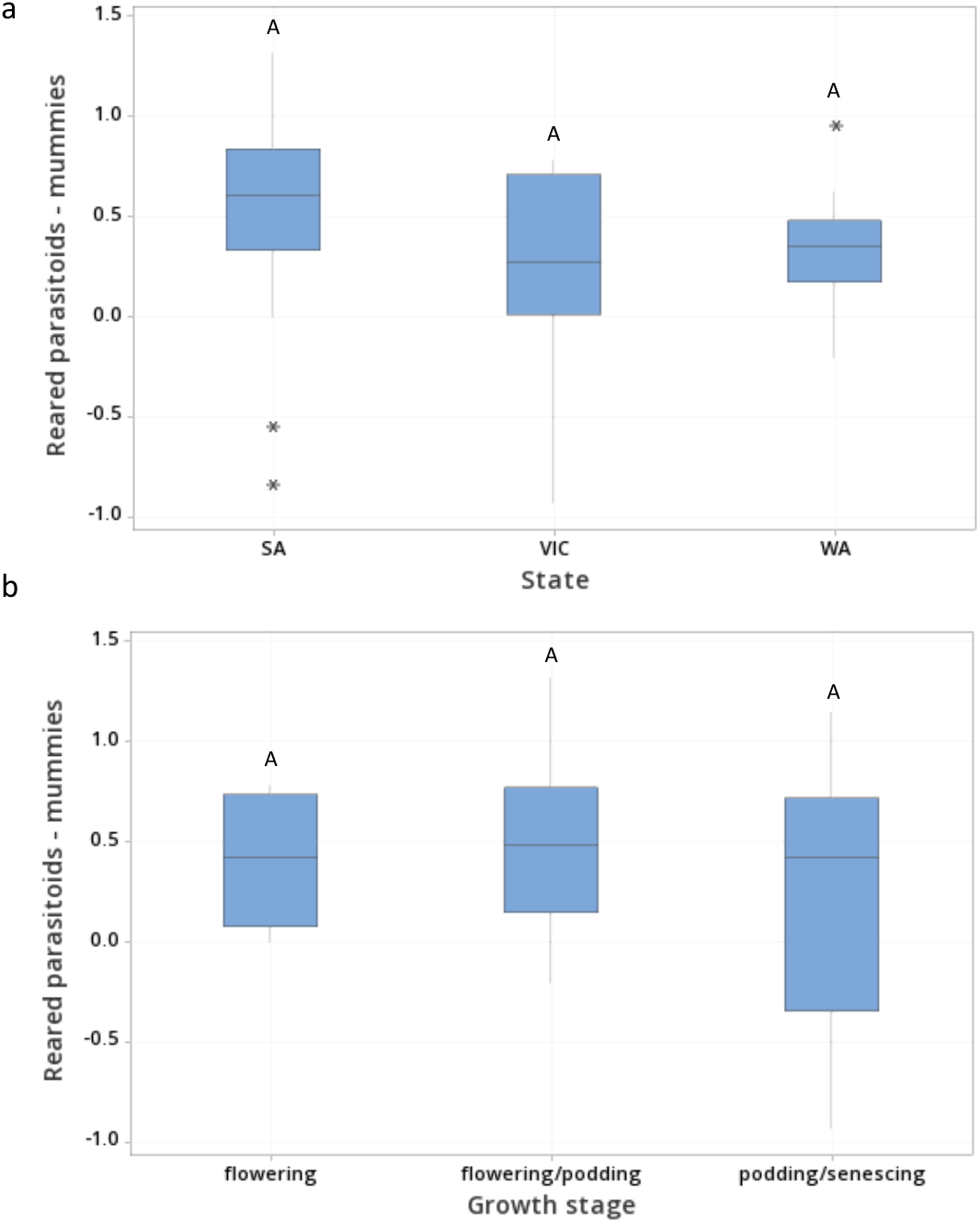
Box plots displaying effects of a) states and b) crop growth stage on the difference between the number of parasitoids reared and the mummies collected [Outliers are indicated by single asterisks].

### 3.3. Correlations between parasitism and aphid abundance

We tested at a field level whether there were associations between aphid, mummy, and parasitoid abundance in addition to proportions of mummies and parasitoids reared. Mummy counts were negatively correlated with *M. persicae* counts during the flowering/podding stage (Spearman; n=14, r=-0.535, p=0.049) but were not correlated during the podding/senescing stage (Spearman; n=16, r=-0.050, p=0.854). Reared parasitoid numbers were not correlated with *M. persicae* counts during the flowering/podding stage (Spearman; n=17, r=0.336, p=0.187), yet were strongly positively correlated during the podding/senescing stage (Spearman; n=15, r=0.845, **p<0.001**) (Fig. S2). Reared parasitoid numbers positively correlated with mummy counts during the flowering/podding stage (Spearman; n=17, r=0.601, p=**0.011**), yet were not correlated during the podding/senescing stage (Spearman; n=15, r=0.350, p=0.201) (Fig. S3).

Proportions of field mummies were not correlated with *M. persicae* abundance during the flowering/podding stage (Spearman; n=21, r=-0.226, p=0.324, but were negatively correlated during the podding/senescing stage (Spearman; n=16, r=-0.888, p**<0.001**). Proportions of reared parasitoids were not correlated with *M. persicae* abundance during the flowering/podding stage (Spearman; n=17, r=-0.083, p=0.750) or the podding/senescing stage (Spearman; n=13, r=0.440, p=0.133). An increase in aphid abundance at the end of the season therefore led to a decrease in observed parasitism but not actual parasitism. Proportions of mummies and proportions of reared parasitoids were highly correlated during the flowering/podding stage (Spearman; n=17, r=0.640, p**=0.006**) however were not correlated during the podding/senescing stage (Spearman; n=13, r=-0.335, p=0.263). These analyses show how parasitism rates in fields can be independent of aphid counts but also how mummification rates may give a different picture on the control provided by parasitoids in a field.

### 3.4. Myzus persicae *abundance and parasitism spatially within a field*

Although our sampling in the field did not proceed perpendicularly from the field edge but proceeded in a zigzag fashion, the distance from the field edge varied suitably to consider distance effects within this context. Only the first eight sampling points (∼210 m moved from the field edge) were used in the analysis as these sampling distances were repeated each visit, regardless of aphid presence/absence. Although there was a state and field effect on abundance of *M. persicae*, there was no effect of distance from the field edge (GLM: state MS=61.436, F_3,19_ =207.53; p**<0.001**; field MS=3.125, F_19,160_ =10.56; p**<0.001**; distance from field edge MS=1.120, F_1,160_=3.78, p=0.054, error MS=0.296) (Fig. S4a). Again, although there was a state and field effect, the proportion of mummies sampled from *M. persicae* was not affected by distance from the field edge (GLM: state MS=3.087, F_3,19_ =5.23; p=**0.002**; field MS=3.396, F_19,142_ =5.75; p**<0.001**; distance from field edge MS=0.387, F_1,142_=0.65, p=0.420, error MS=0.591) (Fig. S4b). The proportion of parasitoids reared was also affected by state and field but not affected by distance from the field edge (GLM: state MS=7.443, F_2,14_ =8.69; p**<0.001**; field MS=3.647, F_14,99_ =4.26; p**<0.001**; distance from field edge MS=0.295, F_1,99_=0.34, p=0.558) (Fig. S4c). The data therefore suggest a relatively even rate of mummification and parasitism across these fields.

### 3.5. Aphid-derived versus mummy-derived parasitoids

The field mummies, of which 370 were collected from SA, VIC and WA, produced 280 parasitoids, with a successful rearing rate of 76 %. Of the 7668 non-mummified *M. persicae* observed in SA, VIC and WA during this study, 708 (9 %) became mummies and produced parasitoids within the laboratory. The aphids that became mummies but did not produce parasitoids were not analysed. Of the parasitoids reared, those reared from field aphids constituted 49 % of total parasitoids reared in WA, 33 % in VIC, and 83 % in SA. On a field level, the number of parasitoids reared from field aphids strongly positively correlated with the number of parasitoids reared from field mummies (Spearman; n=43, r=0.637, p**<0.001**). The difference between the number of parasitoids reared from field mummies and those reared from field aphids (number of parasitoids from field mummies subtracted by parasitoids from field aphids) was not significantly different across field, crop growth stage, or crop stress, but was different across states (GLM: state MS=8.275, F_2,14_=7.94, p=**0.002**; field MS=0.615, F_14,23_=0.59, p=0.846; crop growth stage MS=2.049, F_2,23_=1.97, p=0.163; crop stress MS=2.021, F_1,23_=1.94, p=0.177; error MS=1.042). The difference between the number of parasitoids reared from field mummies and from field aphids per field was significantly greater in SA than in VIC (Fig. 5).

**Figure 5:**
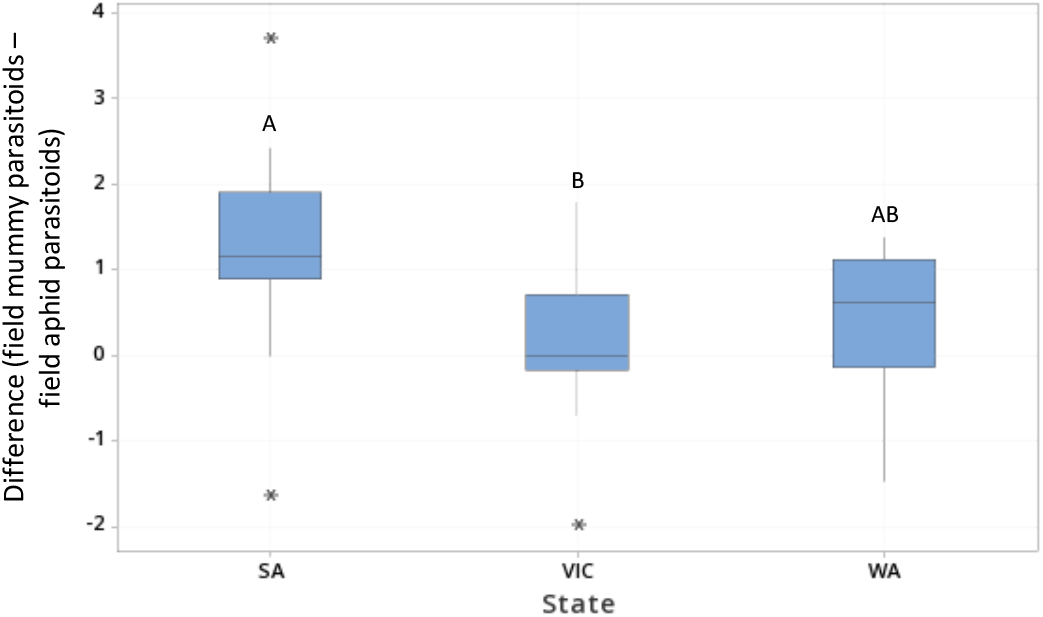
Box plot displaying effects of state on the difference between the number of parasitoids reared from field aphids and those reared from field mummies (number of parasitoids from field mummies subtracted by parasitoids from field aphids) per field. [Outliers are indicated by single asterisks. Different letters indicate significant differences between growth stages in posthoc tests].

### 3.6. Parasitoid community

Primary parasitoids constituted 98 % of all parasitoids reared, hyperparasitoids 2 %, and mummy parasitoids 0.08 %. Of the primary parasitoids, 73 % of those reared were *D. rapae*, 10 % *Aphidius ervi*, 9 % *Aphidius colemani*, with the other species each constituting <5 % of the total (Fig. S5). No Aphelinidae were reared during this experiment. Further details of parasitoid composition by state are given in the supplementary material. Total parasitoid species composition was not different across fields (MRPP, A=0.009, p=0.293) or states (MRPP, A=0.009, p=0.264), nor did it differ with crop stress (MRPP, A=0.009, p=0.300). Crop growth stage, however, was found to have a significant effect on parasitoid species composition (MRPP, A=0.239, p=**0.001**), with there being a greater variety of parasitoid species as plant growth progressed (Fig. 6).

**Figure 6:**
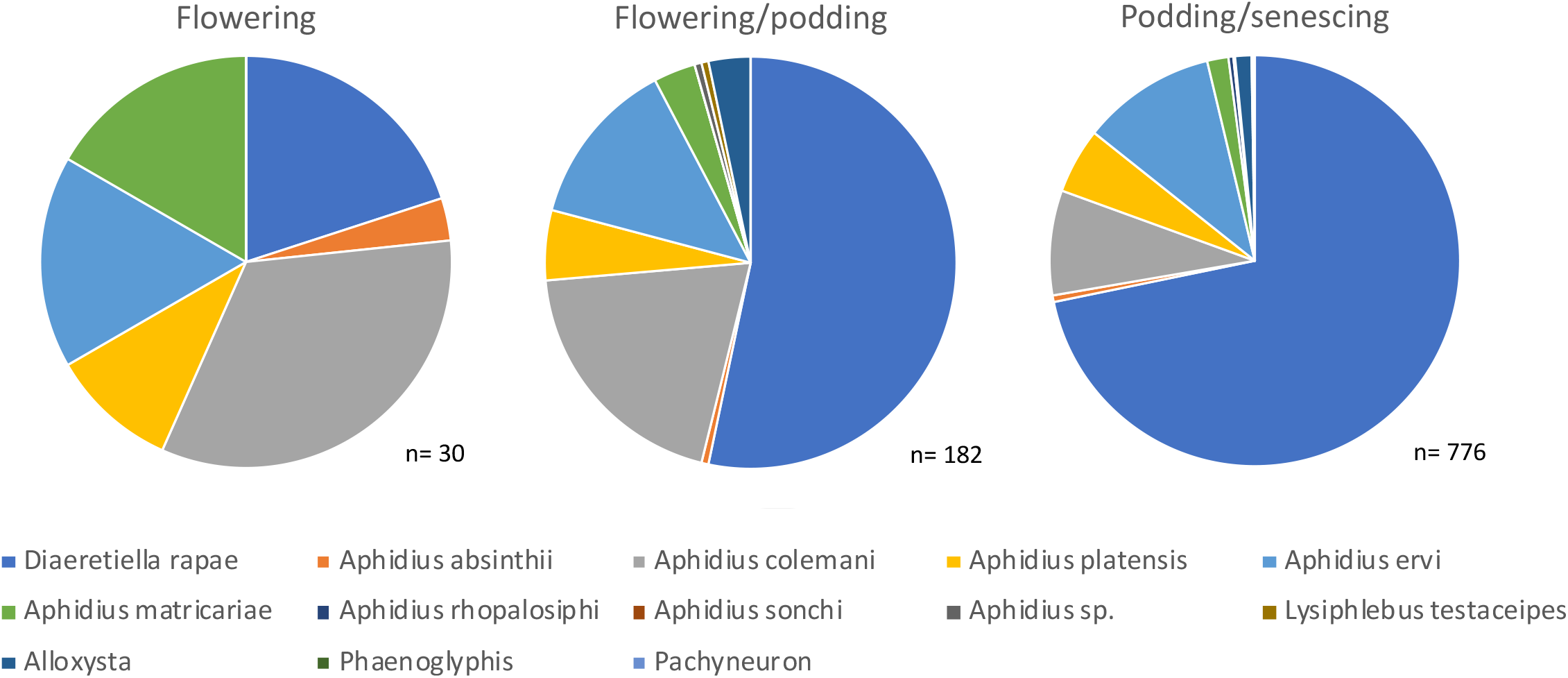
Parasitoid composition within canola crops during the different crop growth stages. Note sample sizes below pie charts.

## 4. Discussion

### 4.1. Parasitism rates

Aphids were detected in August in VIC and July in the other states. Although possibly present in low numbers earlier prior to this, we considered this their first successful colonisation, most likely due to the reduction in efficacy of the canola seed treatments. As expected, due to the timing of aphid arrival, no mummies were found in NSW, SA, and WA until August, and September in VIC. A delay between host and parasitoid emergence is generally observed, particularly in annual crops^24^. At the field level, actual parasitism rate was found to be 2.4 times higher than the observed parasitism rate. The discrepancy was even greater in SA, with actual parasitism rate 4.1 times higher than observed parasitism rate. At the sampling point level, actual parasitism rates were higher for 83% of the points sampled, suggesting that, particularly at this finer scale, observed parasitism usually provides an underestimate of actual parasitism rates. However, due to differences among fields it is difficult to make generalizations, likely due to factors such as landscape composition, configuration, and complexity, and the availability of alternative hosts, and noted in other studies worldwide^25,26^.

Estimates of parasitism levels based on mummy counts (observed parasitism) only relate to numbers of parasitoids at pre-pupal and pupal developmental stages^8^. Around 9% of field-aphids became mummies and successfully reared parasitoids in the laboratory, with the proportion of parasitoids reared from field-aphids highest in SA. It must be noted, however, that rearing parasitoids within a laboratory enables control of climatic variables that could fluctuate in the field, thereby causing failed emergence and parasitoid mortality under more realistic conditions^27^. The difference between the number of parasitoids reared from field mummies than field aphids being significantly greater per field in SA could be due to more sampling occurring during the podding/senescing crop growth stage in this state (at 41%), compared to the other crop growth stages. VIC particularly, had a lot of sampling undertaken during flowering, most likely due to a slower growth rate due to less favourable conditions. This could also explain the lower aphid counts in this state, with aphids being attracted to the yellow colour of the flowering canola and subsequently building in numbers^3^. Any variation in actual parasitism and observed parasitism might therefore be attributed to crop growth stage.

### 4.2. Variation in rates during the canola growing season

Although *M. persicae* abundance and the proportion of alates were not affected by crop growth stage, the proportion of mummies sampled was significantly higher during the podding/senescing growth stage than during earlier crop growth stages, likely due to the delayed presence of parasitoids in the field. The reason for the lack of crop growth stage effects on *M. persicae* abundance could be due to a number of overriding contributing factors that affect aphid population growth, such as elevation, temperature, and/or dew point^28^, in addition to the presence of natural enemies. Interestingly, stress was not found to affect *M. persicae* abundance, or proportions of alates, mummies, or parasitoids reared. It has been said that the formation of aphids’ wings (and in turn dispersal) is indirectly linked to plant stress, but directly to overcrowding, which in turn causes deterioration of the host plant^29^. Plants were identified as ‘stressed’ based on their appearance, which could have been caused by any number of factors, including but not limited to, aphid damage, environment (i.e., moisture or heat stress), or other arthropods (i.e., mite stress); many of which may not be relevant to aphids or parasitoids.

Field parasitism was only observed from August and September onwards, and occurred mostly between the flowering and podding & senescing stages, with parasitoids peaking during flowering/podding and podding & senescing stages. *Myzus persicae* populations and their parasitism vary between years^30^, with the aphid occurring early on in the crop when there is a substantial green bridge (plant material, (i.e., weeds and volunteer plants) surviving over-winter between cropping seasons, that can act as hosts for pests and beneficials)^31^. Further sampling should be undertaken across other years to compare findings.

Although other studies have found that as the season progresses, parasitoids are also more likely to die or diapause within developing mummies^8^, the proportion of reared parasitoids increased as crop growth stage progressed in both SA and WA, but not in VIC. This could be due to the later arrival of aphids in VIC than in the other states, and the subsequent lower proportion of mummies collected. This knock-on effect could be attributed to more fields being sampled during the flowering growth stage and fewer during the podding/senescing stage. The proportion of reared parasitoids in VIC was highest during the flowering/podding stage. Although the flowering/podding crop growth stage in SA and WA spanned the months of August and September, in VIC some of the sampled crops were still flowering/podding in November (coinciding with podding/senescing in the other states). At this time temperatures are higher than earlier in the growing season (∼1.5x higher than in June). The time taken for mummies to form was found to be inversely correlated with the temperature in the range of 10-25°C for *A. colemani* and *A. matricariae*, for example^32^, suggesting the summer month temperature of November is more suited for aphidiine developmental time. These results suggest that the increase in proportion of reared parasitoids is probably more influenced by temperature and seasonal differences rather than the crop growth stage.

The correlation we observed at the field level between *M. persicae* counts and reared parasitoid counts suggests that, when scouting a field to determine parasitism, *M. persicae* numbers alone can be a good indication of the number of parasitoids subsequently reared during the latter growth stages. The proportion of parasitoids reared, however, cannot be accurately determined using this method as there appears to be only a weak correlation between these two variables. Moreover, the data suggest that the relationships between observations in the field and actual parasitism may vary during the season.

### 4.3. Composition of aphid parasitoids

The parasitoids comprised mostly of primary parasitoids, with *D. rapae* the dominant species. Secondary parasitoids were very low throughout the season. It is important to assess the presence of hyperparasitoids (or mummy parasitoids) which could affect the long-term ability of primary parasitoids to control *M. persicae* populations. Parasitoid species composition was not affected by state, field or crop stress, but was affected by crop growth stage, with *D. rapae* becoming more dominant as crop growth stage progressed. This could reflect warmer temperatures during the latter growth stages, as *D. rapae* developmental time decreases more rapidly than that of other aphidiines, such as *A. matricariae*, as temperature increases^33^. *Diaeretiella rapae* has been found to be very tolerant of drought conditions (in the absence of increased temperature), continuing or even increasing aphid parasitism^34^. This could explain the dominance of this species in southern Australia given that 2019 was a drought-affected year.

Additionally, *D. rapae* has a very broad range of host aphids and plants compared to other aphidiines, and so has the ability to host swap and build up populations outside of the growing season^35^. More *D. rapae* were reared in WA, likely due to higher abundance of *B. brassicae* in this state, as this aphid species is reputedly the preferred host of *D. rapae*^36^. It has been recorded that *D. rapae* does not respond to chemical attractants from aphids, as increasing aphid densities did not affect the arrival time of this parasitoid^37^ and may respond instead to volatile compounds from cruciferous crops^38,39,40^. *Diaeretiella rapae* orientates towards mustard oils produced by crucifers, causing it to attack aphids on this plant type^41^. This ability to respond to plant volatiles potentially increases its efficacy, as volatile chemicals released by plants can be important when parasitoids locate hosts^42^.

The difference between observed and actual parasitism varied little between primary and secondary parasitoids, and between *D. rapae* and other primary parasitoids, so that parasitoid species composition had no discernible effect on observed and actual parasitism rates. The difference in observed and actual parasitism rate also did not vary across state, field, crop growth stages or crop stress, suggesting there are no clear trends against these factors.

### 4.5. Conclusion

Observed and actual parasitism rates of *M. persicae* can vary considerably, due to a number of variables ranging from climatic factors to faunal composition. Parasitoid species, however, appeared to have no effect on the difference between observed and actual parasitism rates. On a state, field and sampling point level, actual parasitism was usually higher than that observed in the field. Average actual parasitism rate was over double that observed, and even more pronounced in SA, with mummy counts usually an underestimate of actual parasitism rates. This provides an incentive to incorporate natural enemies into pest management programs. Consequently, mummy counts alone do not provide a clear representation of parasitism within the field and can vary across geographic areas or within growing seasons.

## Acknowledgements

We would like to acknowledge Andy Hulthen at CSIRO for his creation of the mobile software application. Furthermore, we would like to thank the growers and agronomists who assisted with this project and/or provided access to their land. We further extend our gratitude to Sarina Macfadyen for her supervision, Katie Robinson and Moshe Jasper for their assistance with genetic work and training, Vincent Chea for his GIS expertise, and Erica Marshall for her help with R software. This research initiative is a GRDC investment that seeks to deliver new knowledge to improve the timing of pest management decisions in grain crops to grain growers: CSE00059. This work was further supported by the Albert Shimmins Fund.

## Conflict of Interest Declaration

The authors declare no conflict of interest.

## Notes

### Competing Interest Statement

The authors have declared no competing interest.

