## Supplementary material for "Is what you see what you get? The relationship between field observed and actual aphid parasitism rates in canola crops": S

*Table S1:* Collection data acquired during field sampling ('->' sign indicates a subcategory (drop down list) that appears once the previous row has a positive response).

| Data type | Description | Categories (explained within text) |
|---|---|---|
| Collector | Name of person collecting data | N/A |
| Field ID | Unique field identifier (name of paddock) | N/A |
| Growth stage | Crop information; crop growth stage | Seedling; vegetative; early flowering; flowering and podding; podding only; senescing; other |
| Condition | Crop information; plant stressor type | No visible stress; moisture stress; mite stress; stunted; patchy; other |
| -> Condition (other) | Crop information; plant stressor type, if not listed above | N/A |
| Wasp presence | Presence of aphid parasitoids on yellow sticky traps (data not used within this paper) | Present; absent |
| Crop notes | Any additional information found to be necessary | N/A |
| Longitude/ latitude | Sample sites; G.P.S. coordinates (updated automatically) | N/A |
| Altitude | The height of the site (in metres) in relation to sea level | N/A |
| Accuracy | Accuracy of GPS data (in metres) | N/A |
| Pest presence | Presence of aphid pests. (Up to 8 points where aphids are present should be sampled, or maximum of 24 sampling points) | Yes; no |
| -> Pests | Species of pest aphid present | Green peach aphid; cabbage aphid; turnip aphid; other |
| -> *M. persicae* location | If *M. persicae* present, where were they present on a plant? | Lower leaves; middle leaves; upper leaves; racemes; other |
| -> *M. persicae* location (other) | Location of *M. persicae*, if present and not listed above | N/A |
| -> *M. persicae* counts | If *M. persicae* present, how many of each form were there? [Drop down menu for separate counts] | *M. persicae* winged [alates]; *M. persicae* adults [apterae]; *M. persicae* nymphs; mummies closed; mummies open |
| -> Alternate collection | When *M. persicae* not present but another species of pest aphid is, this should be collected | Yes; no |
| Natural enemy presence | Presence of natural enemies associated with aphids | Yes; no |
| -> Natural enemies | Type of natural enemy present (both arthropod type and form) | Lacewing larvae; lacewing adults; ladybird larvae; ladybird adults; hoverfly larvae; hoverfly adults; parasitoids; predatory beetles; predatory bugs; spiders; pathogenic fungi; other |
| -> Natural enemies (other) | If natural enemies were present that were not listed above | N/A |
| Sample ID | Barcodes assigned to each site, to identify collections taken into the laboratory for rearing | N/A |

*Table S2:* Processing data required during laboratory rearing process ('->' sign indicates a subcategory (drop down list) that appears once the previous row has a positive response)

| Data type | Description | Categories (explained within text) |
|---|---|---|
| Collector | Name of person collecting data | N/A |
| Processing stream | Aphid form parasitoid was reared from | Mummies; un-parasitised |
| -> Sample ID | Barcodes assigned to each site, to identify collections taken into the laboratory for rearing | N/A |
| -> Emerged wasps | Number of parasitoids emerged from either mummies or seemingly un-parasitised aphids | N/A |
| -> Empty mummy cases | Number of empty mummy cases pertaining to the emerged parasitoids | N/A |
| -> Unsuccessful rearings / Un-mummified aphids | Depending on the 'processing stream' selected, 'unsuccessful rearings' is linked to the 'mummies' stream and 'un-mummified aphids' is linked to the 'un-parasitised' stream, but both pertain to aphids that did not rear parasitoids | N/A |

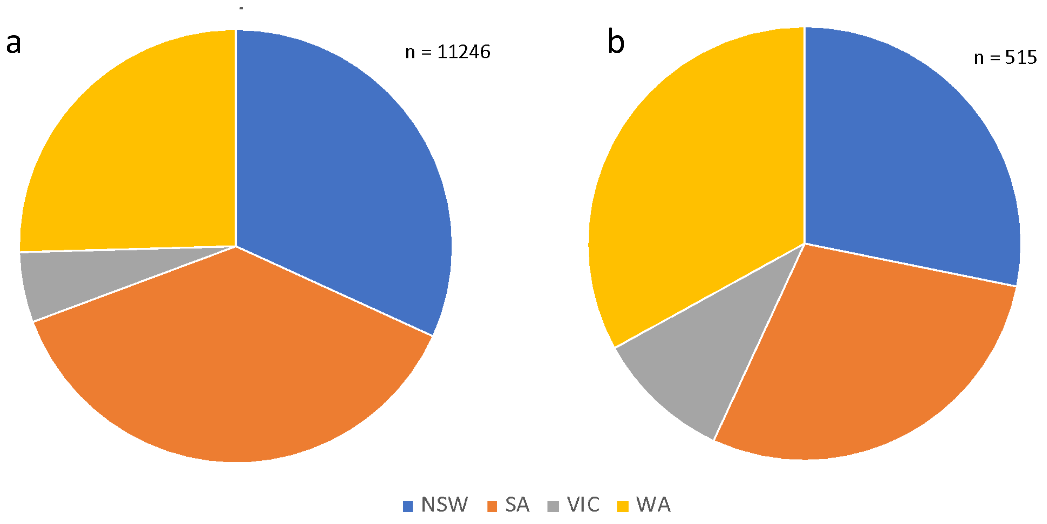


*Figure S1:* Proportion of a) non-mummified and b) mummified *M. persicae* for each state when summed across sites and collections.

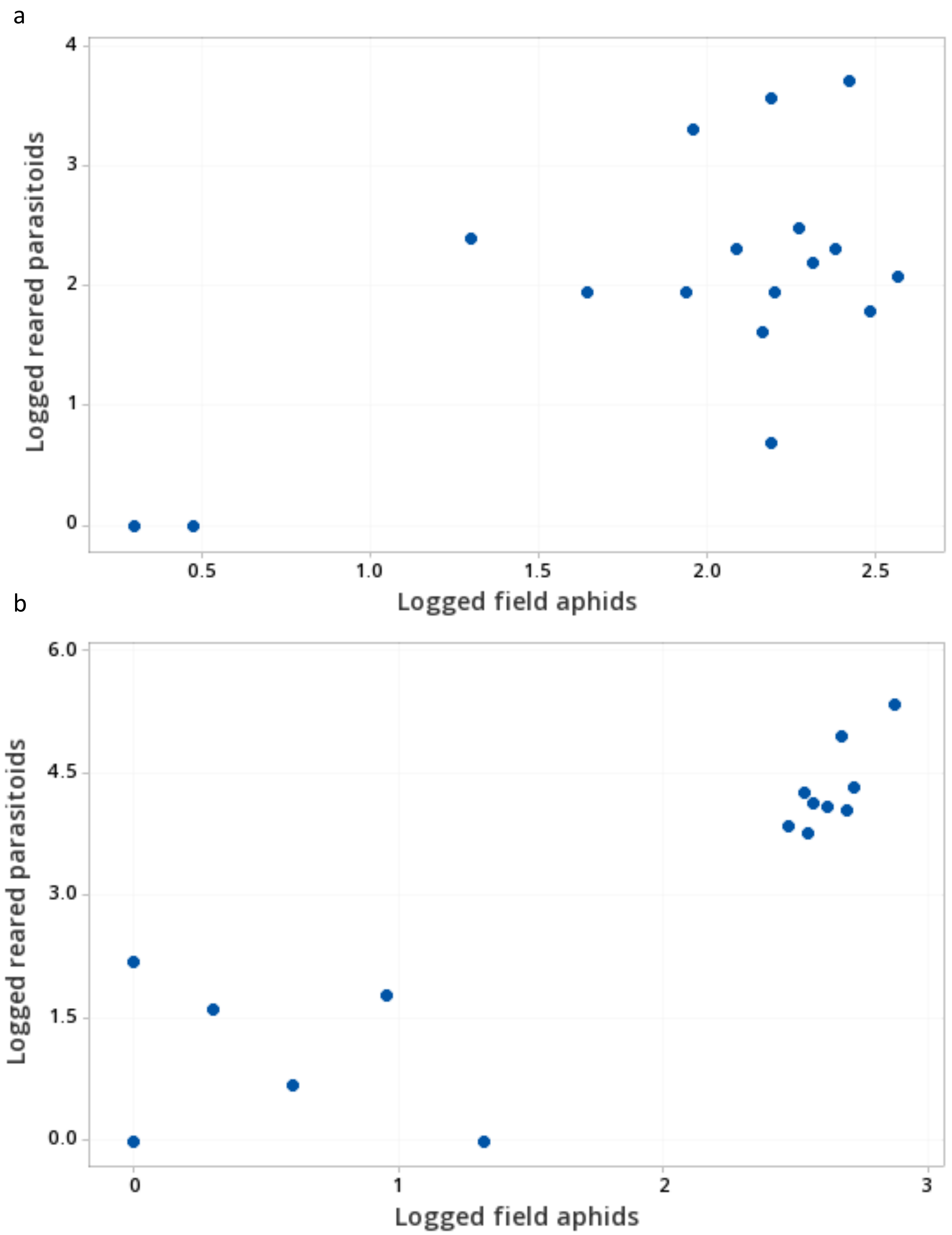


*Figure S2:* Correlation of reared parasitoid numbers with field aphid counts during the a) flowering/podding stage and the b) podding/senescing stage [Counts as (ln(x+1))].

a

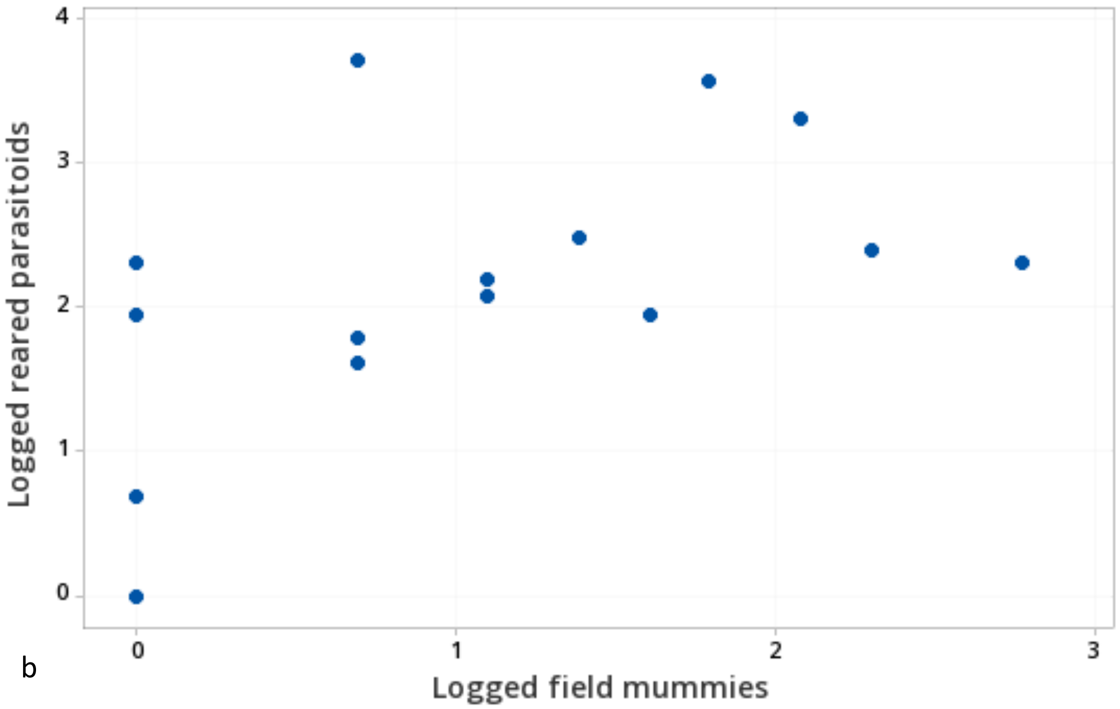


b

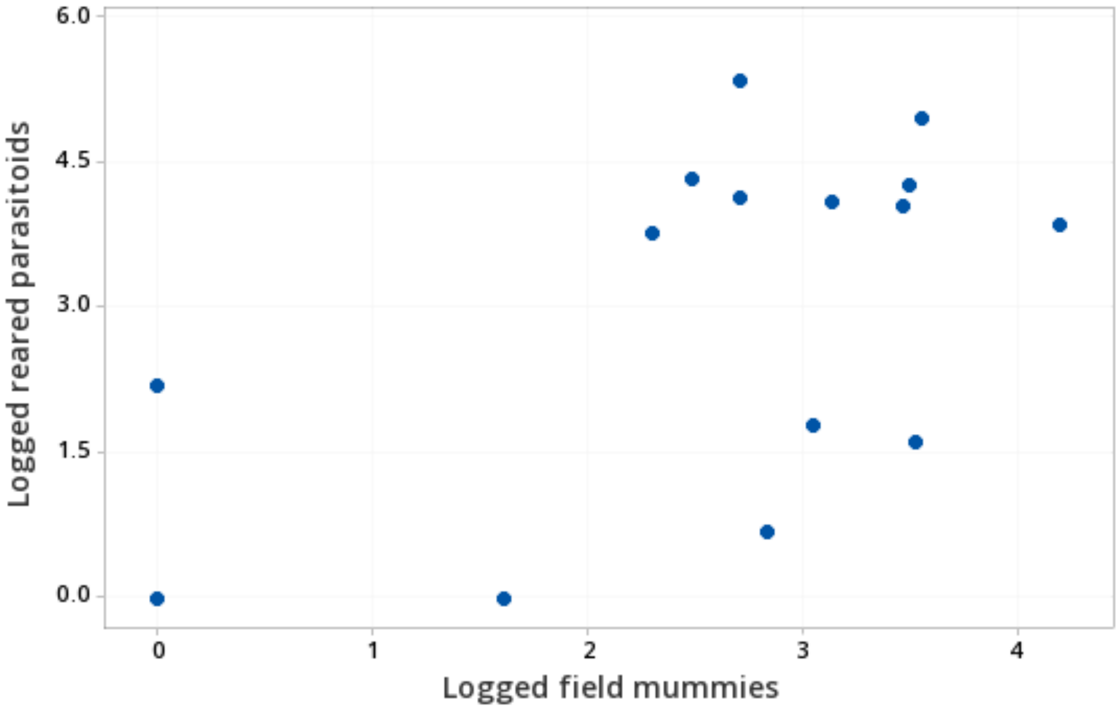


*Figure S3:* Correlation of reared parasitoid numbers with field mummy counts during the a) flowering/podding stage and the b) podding/senescing stage [Counts as (ln(x+1))].

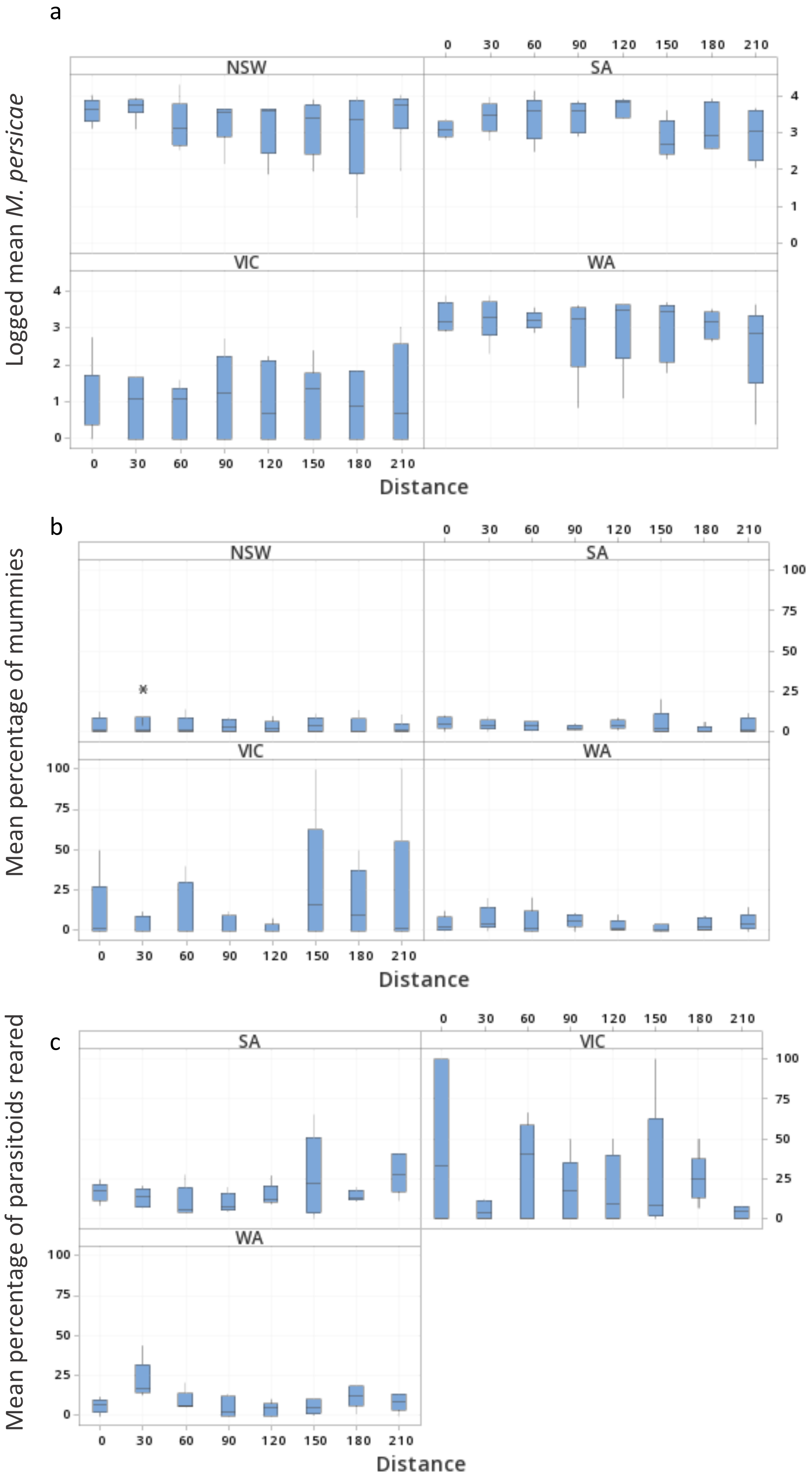


*Figure S4*: Effect of distance from paddock edge on a) logged *M. persicae* abundance (ln(x+1)), and percentage of b) mummies and c) reared parasitoids.

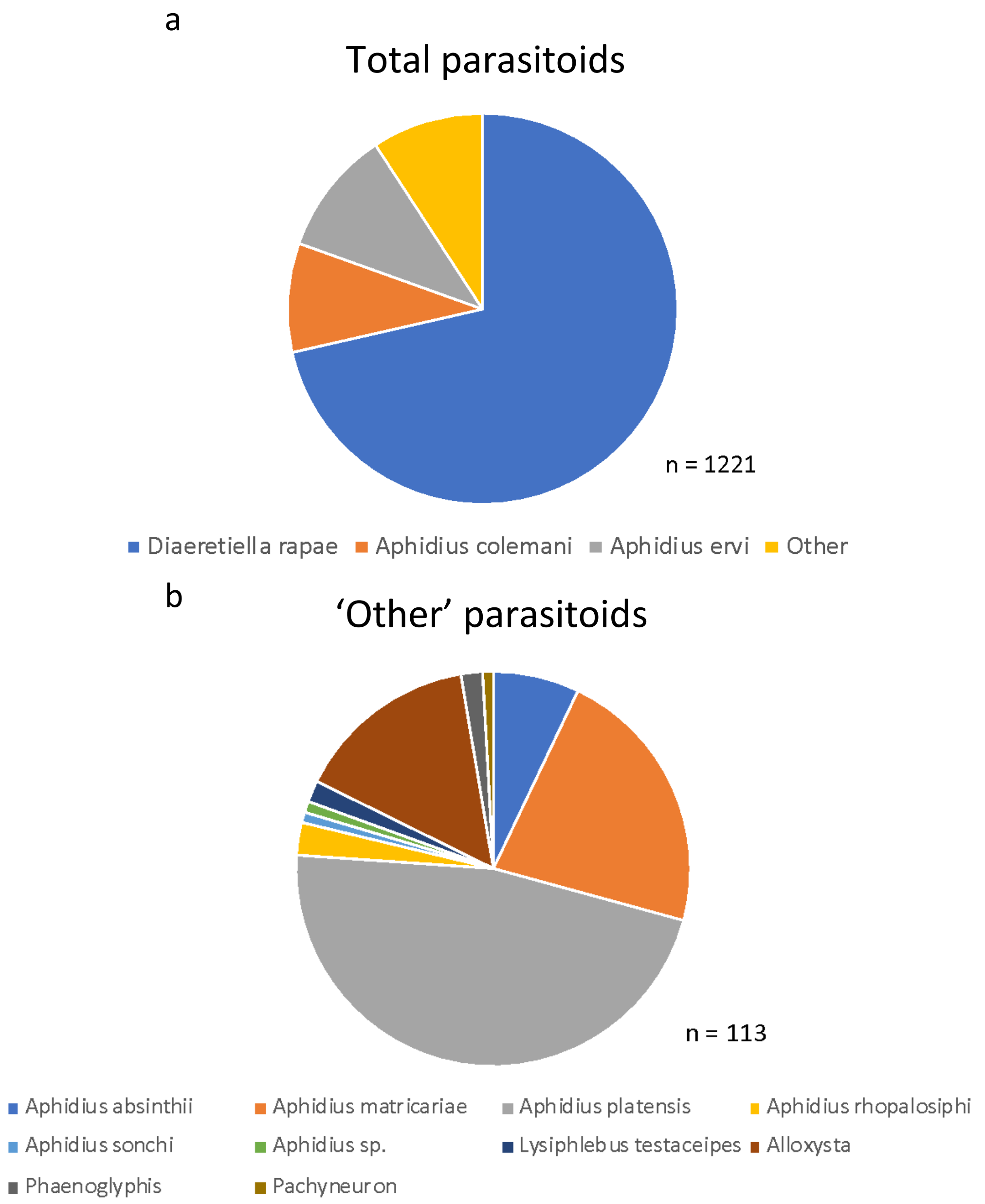


*Figure S5:* Parasitoid species composition of parasitoids reared from *M. persicae* from within Australian canola crops.

In VIC, primary parasitoids constituted 92 % of all parasitoids reared compared to 99 % in NSW, SA and WA. Of the primary parasitoids, *D. rapae* was always the predominant parasitoid, constituting 92 % in NSW, 62 % in SA, 69 % in VIC, and 81 % in WA. Hyperparasitoids constituted 1 % in NSW, SA and WA, and 7 % in VIC. Only one mummy parasitoid was reared: a *Pachyneuron* sp. in VIC. The majority of hyperparasitoids comprised *Alloxysta* sp., with the exception of two *Phaenoglyphis* sp. individuals, one collected from NSW and one from WA.

Reared parasitoids from SA, VIC and WA were separated into *Diaeretiella rapae* (M'Intosh), the most commonly reared parasitoid, and non-*D. rapae* species for each sampling point. The number of reared *D. rapae* increased with crop growth stage (Fig. 6), and greater numbers of *D. rapae* were reared in WA than from the other states. The proportion of *D. rapae* and non-*D. rapae* species reared from field-aphids was similar: 27 % of *D. rapae* were reared from field-mummies and 73 % from field-aphids, while 31 % of non-*D. rapae* were produced from field-mummies and 69 % were reared from field-aphids. When analysed at the level of fields, the proportion of *D. rapae* was not significantly different to the other parasitoids when reared from field aphids versus field mummies ($t_{(43)}$=1.91, p=0.070).

Of the primary parasitoids 28 % were reared from field-mummies and 72 % from field-aphids, whereas 67 % of secondary parasitoids were reared from field-mummies and 33 % from field-aphids. The number of secondary parasitoids reared, however, was low (18).
